# Mechanistically explainable AI model for predicting synergistic cancer therapy combinations

**DOI:** 10.1101/2025.08.10.669494

**Authors:** Han Si, Sanyam Kumar, Sneh Lata, Arshad Ahmad, Saurav Ghosh, Karen Stephansen, Deepti Nagarkar, Eda Zhou, Brandon W. Higgs

## Abstract

This study introduces a Large Language Model (LLM)-based framework that combines drug combination data with a knowledge graph to predict synergistic oncology drug combinations with mechanistic insights. Using a retrieval-augmented generation (RAG) approach, over 50,000 *in vitro* drug pair assay results and 1,631 human clinical trial or preclinical test entries were integrated to enhance predictive accuracy and explainability. Validation achieved an F1 score of 0.80, demonstrating the framework’s potential to streamline drug discovery and improve translational strategies in cancer treatment.

---

The complexity and adaptability of cancer make single-agent treatments less effective, as tumors quickly develop resistance^1^. Drug combinations address this by targeting multiple pathways concurrently, reducing resistance and often enhancing effectiveness^2^. For instance, pairing immune checkpoint inhibitors with chemotherapy boosts immune response while targeting tumor cells^3, 4^. Similarly, combining BRAF and MEK inhibitors has improved outcomes in melanoma^5, 6^.

Over the past two decades, *in silico* methods for predicting drug synergy have advanced drug combination discovery, reducing the need for resource-intensive lab experiments^7^. Early models integrated transcriptomic and chemical-genomic data^8^, while initiatives like the NCI-DREAM Challenge spurred models such as DeepSynergy^9, 10^. Recently, transformer-based models trained on large-scale *in vitro* datasets like CancerGPT have further improved predictive accuracy^11^.

Despite advancements, translating animal and cell line model findings to human trials remains challenging, particularly in providing mechanistic explanations for drug combinations^12^. Our study builds on prior methods with a new framework integrating phase 1-3 clinical trial outcomes, *in vitro* screens, and a biological knowledge graph (PrimeKG) to improve mechanistic insights^13^. By combining 1,631 human study results across 723 drug modality combinations (Supplemental Figure 1), and 50,000 *in vitro s*creens into a RAG model^14^ with entity relationships (Figure 1), our approach offers biologically grounded, interpretable predictions, to improve discovery and forward/reverse translational stages of clinical development.

**Figure 1.**
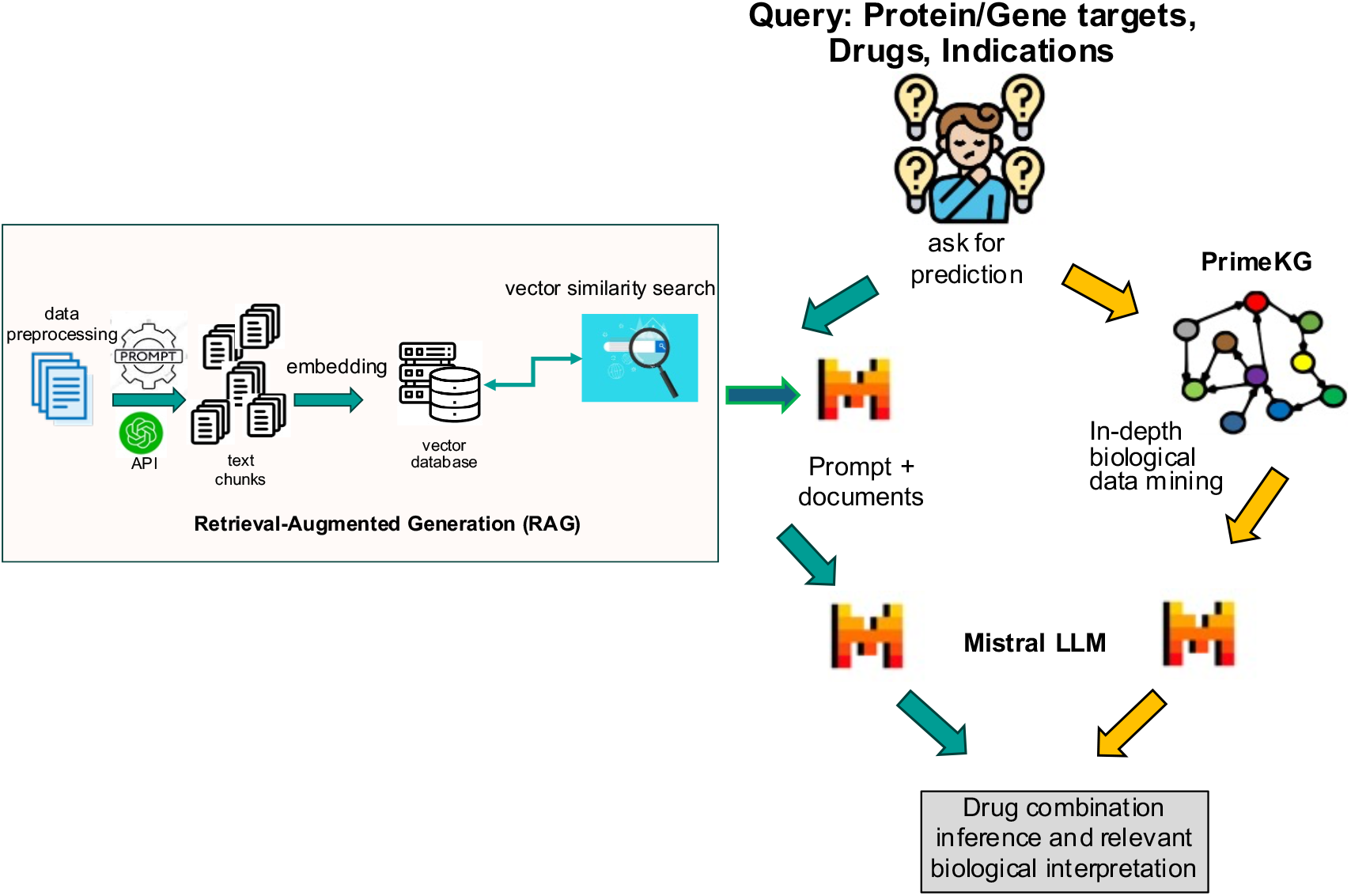
Model architecture illustrating the input data flow as a RAG into a vector database that was queried by the user [Drugs or Gene/Protein targets, Indications] in the LLM and the knowledge graph to provide a prediction result with biological rationale for the drug combination in the specific indication.

A validation set inclusive of drug combinations and tumor types both seen and unseen to the input RAG was manually curated (Figure 2). This dataset consisted of 58 test cases across a diverse set of tumor types, patient genetic subtypes, and oncology drug modalities involving combinations of immuno-oncology (IO) therapies, targeted therapies, antibody-drug conjugates (ADCs), and chemotherapies. Figure 3A illustrates the model accuracy, precision, recall, and F1 score for this validation dataset with test cases seen and unseen to the RAG (overall N=58), seen (N=16), and unseen (N=42). Across all drug modalities, the model showed overall accuracy of 77.6%, with cases seen of 87.5%, and unseen of 73.8%. The F1 scores of overall, seen, and unseen test cases were 84%, 92.3%, and 80%, respectively. For F1 scores across drug modalities (Figure 3B), ADC agents in combination with other therapies performed best (100%), which tied with performance of targeted therapies combined with either other targeted or chemotherapy agents (100%), while IO-IO combinations performed the worst (66.7%). To illustrate the mechanistic explainability of the predictions, a few examples unseen to the model are provided below with additional examples in Supplementary Data.

**Figure 2.**
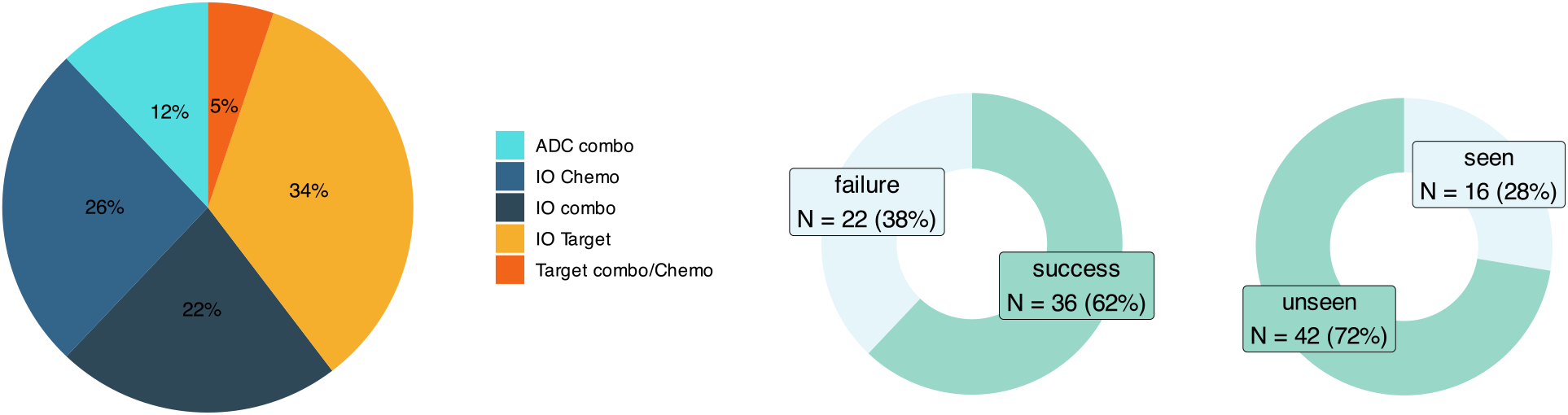
Distribution of manually curated validation set. Graphs show breakdown of drug combination modalities, success/failure status of clinical trial, seen/unseen to the RAG

**Figure 3.**
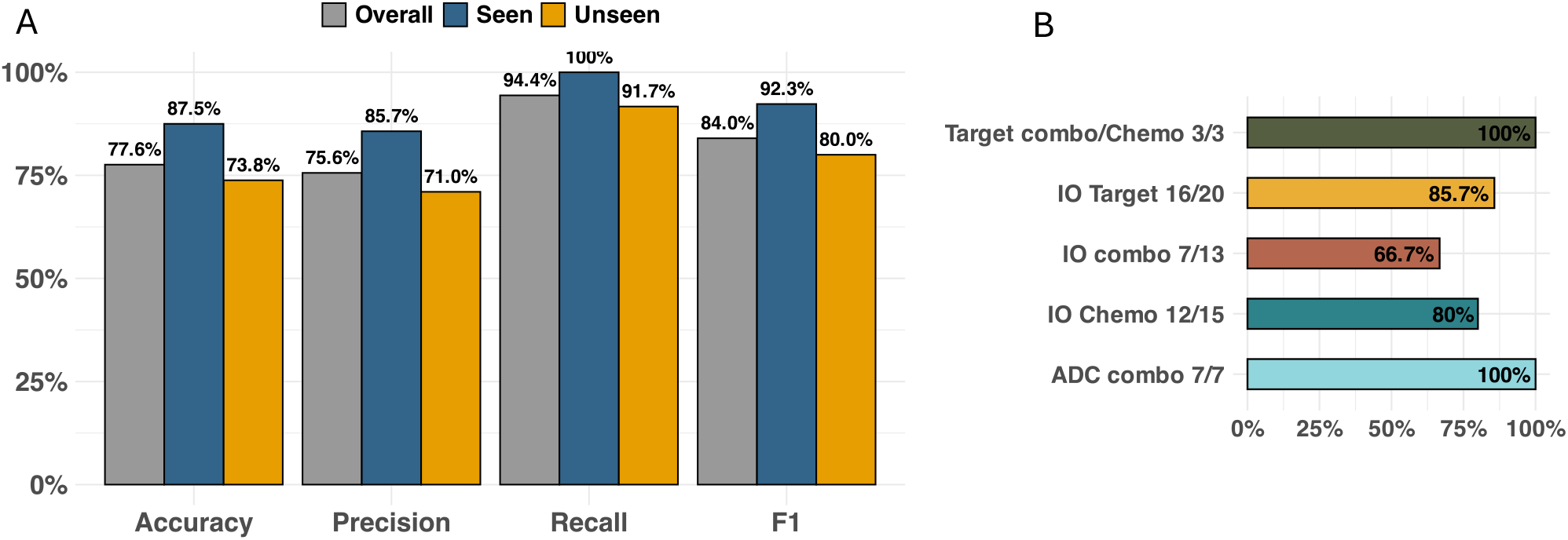
Model performance across all drug modalities in (A) overall, RAG seen, and RAG unseen cases and (B) F1 scores separated by drug modality combination in all cases.

The model successfully predicted the combination of atezolizumab (anti-PD-L1) and cobimetinib (MEK1/2 inhibitor) to not be effective in metastatic colorectal cancer (CRC), consistent with the phase 3 IMblaze370 trial, where the combination failed to improve overall survival compared to regorafenib^15^. Beyond accuracy, the model provided detailed mechanistic reasoning, abbreviated here: “*Atezolizumab is an anti-PD-L1 monoclonal antibody that blocks this interaction, thereby restoring T-cell function and enhancing the immune response against cancer cells. Cobimetinib, on the other hand, is a MEK1/2 inhibitor that blocks the MAPK signaling pathway, which is involved in cell growth and survival. While both drugs have shown activity in various cancers, there is no definitive data on their synergy in locally advanced or metastatic CRC. In fact, there is evidence suggesting that MEK inhibitors may decrease the efficacy of PD-1 blockade by increasing the expression of PD-L1 in some cases. Therefore, it is unlikely that the combination of atezolizumab and cobimetinib would have a synergistic effect in treating locally advanced or metastatic CRC*.” This prediction aligned with clinical outcome, highlighting the model’s mechanistic explanation. MEK inhibition can upregulate PD-L1^16, 17^, potentially increasing targets for anti-PD(L)1 therapy, however, this may also promote an immunosuppressive environment, thus explaining the lack of synergy in CRC.

A true positive prediction involved combining mirvetuximab soravtansine (anti-FOLR1 + tubulin inhibitor) with bevacizumab (anti-VEGFA) in platinum-resistant ovarian cancer. An abbreviated excerpt of the model’s rationale highlighted complementary mechanisms: “…*distinct mechanisms of actions targeting different molecular pathways involved in cancer progression…By targeting FOLR1, mirvetuximab soravtansine can disrupt the folate metabolism pathway and inhibit cell division, making the cancer cells more susceptible to chemotherapy…*.*By inhibiting VEGF, bevacizumab can prevent the formation of new blood vessels, starving the tumor of nutrients and oxygen, and enhancing the efficacy of chemotherapy. The combination of these two drugs can lead to a synergistic effect by targeting different molecular pathways and enhancing the overall therapeutic effect against platinum-resistant ovarian cancer*”. This synergy prediction demonstrates its potential to guide treatment in resistant tumors.

While the model performed well overall, IO-IO combinations posed challenges, often resulting in false positives where the model predicted success, but clinical trials did not. For instance, pembrolizumab (anti-PD1) and vibostolimab (anti-TIGIT), with or without docetaxel, were predicted to be effective in metastatic NSCLC. The model rationale suggested that dual PD-1 and TIGIT blockade would reactivate T cells, while docetaxel would create a more immunogenic environment. However, in the phase 2 KeyVibe-002 trial, this combination failed to improve progression-free survival^18^. To reduce such mispredictions, incorporating more examples of failed trials is essential.

Our approach significantly improves upon previous models by not only expanding the input training content on drug combinations, which to date have relied heavily on *in vitro* drug screens, but also by providing either successful/failed studies in a clinical setting and an interpretable framework for both predictive power and biological insight. Unlike generic LLMs such as ChatGPT-4o, which often generated inaccurate biological explanations (data not shown), our model minimized “hallucinations.” Direct comparisons to ChatGPT-4o are challenging, as it lacks controls to prevent data leakage—where unseen drug combinations could still inform its responses. In contrast, our framework deliberately excluded test cases in the RAG to measure predictive accuracy accurately, a feature that general LLMs with full data access cannot replicate.

Despite these promising results, limitations include lack of diversity in model training such as drug modalities (e.g. CAR-T, cancer vaccines), ADC payloads (e.g. topoisomerase I inhibitor, microtubule inhibitor), dosing considerations, and biases from published successful trials. Expanding these factors will improve accuracy. Despite these considerations, such predictive frameworks can ultimately accelerate the discovery of synergistic therapies, thereby reducing costly experiments, and improve overall patient outcomes in oncology.

## Methods

The following methods are provided in Supplementary Data.

*Pubmed Data Collection*

*Clinical Trial Data from Citeline Trialtrove*

*In vitro Drug Combination Screening Database Knowledge Graph for Biological Explainability*

### Model Architecture

The model architecture consisted of a hybrid framework integrating an LLM with a knowledge graph (Figure 1). The first step was to implement Retrieval-Augmented Generation (RAG): a corpus of drug treatment data from Trialtrove, publication summaries, and DrugComboDb were embedded into a text-based vector database. This RAG served as the foundation for the retrieval step, ensuring the model had access to high-quality, relevant data. Next was to craft prompts to harness the full potential of the LLM, guiding it to focus on accurate drug combination predictions using the retrieved information from the RAG; each prompt aligned with the specific nuances of drug combinations in the query, ensuring the output was both precise and insightful. The final step utilized the knowledge graph, which enriched the model’s predictions by anchoring on biological and disease context. By linking drug-target interactions to known disease biology, the knowledge graph enhanced interpretability, making the model’s predictions not only mere predictions but also providing biologically informed insights.

### Software and Evaluation Metrics

Software licensed from Lamini (Menlo Park, CA, USA) was used for model implementation. Mistral v0.2 was the LLM in the model architecture. Model performance included standard classification metrics such as precision, recall, F1 score, and accuracy. Predictions were also assessed for biological plausibility based on known mechanisms of action from the literature.

## Author contributions

BWH and HS conceived the study design, BWH was a major contributor in writing the manuscript. HS analyzed and interpreted the data and contributed to drafting manuscript, SK provided clinical trial data and contributed to drafting manuscript, SL compiled trial data, AA and SG provided IT and software support, KS and DN managed the project and budget, EZ provided the programming support. All authors edited and approved the final draft of the manuscript.

## Acknowledgements

This study is sponsored by Genmab.

## Data availability

The datasets used and analyzed during the current study are available from the corresponding author upon request.

