## Supplemental Figure 1, Methods for "Mechanistically explainable AI model for predicting synergistic cancer therapy combinations"

**Supplemental Data**

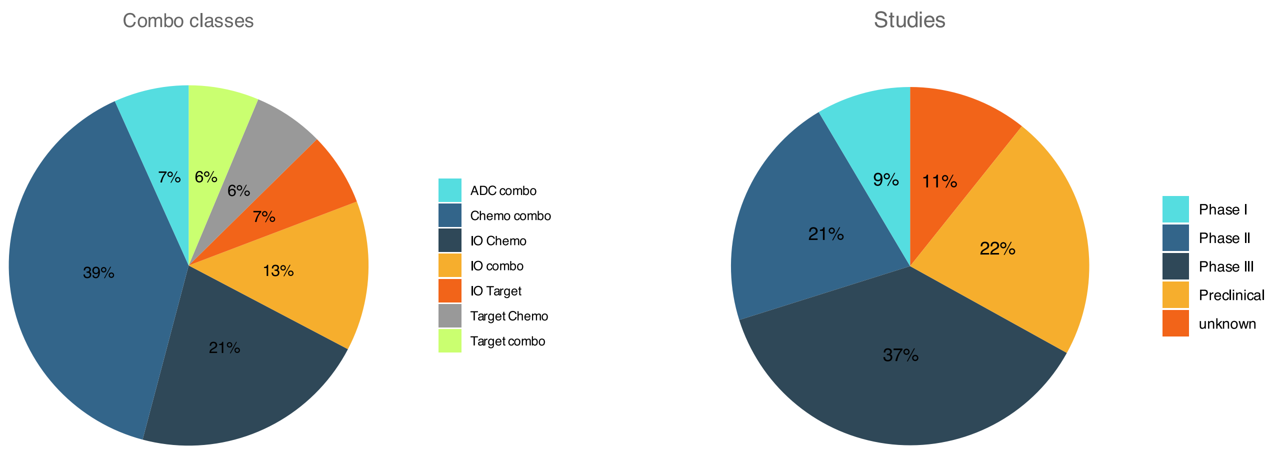


Supplemental Figure 1. Distribution of drug modality combinations (left, N=728) and total drug studies (right, N=1,631) used in the model.

### Supplemental Methods

*Pubmed Data Collection*

Relevant publications were accessed through the NCBI Entrez E-utilities API, dynamically generating queries using drug names and journal titles to target articles on specific drug combinations. Retrieved data were processed using BeautifulSoup. OpenAI GPT-4 extracted biomedical data from abstracts and full-texts. Tailored prompts facilitated three types of extraction: (1) abstract summarization, highlighting tumor type and drug efficacy, (2) drug target identification, and (3) tumor type and clinical trial outcomes. Outputs were validated by manual review and cross-referencing with DrugBank to ensure alignment with established biomedical data. Error handling was integrated for API call robustness.

*Clinical Trial Data from Citeline Trialtrove*

Additional phase III clinical trial data were obtained from Citeline Trialtrove, focusing on trial ID, phase, and outcomes. GPT-4 identifed drug targets, categorizing trial outcomes as Positive,

Negative, or Unknown. The total set of unique drug modality combinations and clinical trial numbers are provided in Supplementary Data.

*In vitro Drug Combination Screening Database*
DrugComboDb[1] was downloaded (release date 2019.05.31), inclusive of >50,000 drug combinations across 51 cancer cell lines; synergy scores were categorized using ZIP, HSA, Loewe and BLISS scores (synergy: if ZIP>10 or any other 2 scores >10; antagonism: if ZIP <10 or any other 2 scores <10; otherwise inconclusive). Drug names were merged with CancerDrugsDB to associate protein targets with each drug, and Cancer Cell Line Encyclopedia (CCLE)[2] to associate cell lines with the tumor from which they were derived.

*Knowledge Graph for Biological Explainability*
A biological knowledge graph containing proteins, pathways, diseases, and drugs, (7,080 diseases with 4,050,249 relationships) enhanced model explainability using PrimeKG.

**Additional drug combo synergy prediction results:**

- Drug combo: Encorafenib, Cetuximab, Binimetinib
- Targets: BRAF, EGFR, MAK
- Disease: BRAF **V600E**-mutated metastatic colorectal cancer
- Inferred synergy: Yes
- (True positive)

Explanation of prediction

- The synergistic effect of the drug combination of Encorafenib, Cetuximab, and Binimetinib in treating BRAF V600E-mutated metastatic colorectal cancer can be explained through their mechanisms of action and the interconnectedness of the molecular targets in the RAS/RAF/MAPK signaling pathway.
- Encorafenib is a BRAF inhibitor, which blocks the activity of the BRAF protein, preventing it from promoting the growth of cancer cells. The BRAF V600E mutation leads to the production of a constitutively active BRAF protein, which results in the hyperactivation of the MAPK signaling pathway.
- Binimetinib is a MEK1/2 inhibitor, which blocks the downstream effectors of the BRAF signaling pathway, leading to the suppression of the MAPK signaling pathway. By inhibiting both the upstream BRAF and downstream MEK1/2 enzymes, the drug combination can effectively block the RAS/RAF/MAPK signaling pathway, leading to a more potent inhibition of cancer cell growth than using either drug alone.
- Cetuximab is a monoclonal antibody that binds to the EGFR protein, inhibiting its activity and leading to tumor cell death. However, the efficacy of cetuximab in BRAF V600E-mutated metastatic colorectal cancer is limited due to the activation of the MAPK pathway. By inhibiting the MAPK pathway with Binimetinib, the activity of Cetuximab can be enhanced, leading to a more effective inhibition of cancer cell growth.

- Therefore, the synergistic effect of the drug combination can be attributed to the complementary mechanisms of action of the three drugs in targeting different components of the RAS/RAF/MAPK signaling pathway, leading to a more potent inhibition of cancer cell growth than using any single drug alone

- Drug combo: Bevacizumab, Pembrolizumab
- Targets: VEGFA, PD-1
- Disease: Glioblastoma
- Inferred synergy: No
- (True negative)

Explanation of prediction

- Based on the given knowledge graph, there is no direct evidence of synergy between Pembrolizumab (targeting PD-1) and Bevacizumab (targeting VEGFR) in the treatment of Glioblastoma. While both drugs have been used in the treatment of various types of cancer, including glioblastoma, their mechanisms of action and targets are distinct.
- Pembrolizumab is a monoclonal antibody that blocks the PD-1 protein, which helps to restore the immune system's ability to fight cancer.
- Bevacizumab is also a monoclonal antibody that inhibits the action of VEGF, which plays a role in angiogenesis and tumor growth.
- While there is some evidence that PD-1 and VEGF may interact in certain biological contexts, there is no clear evidence to suggest that the combination of these drugs would result in a synergistic effect in the treatment of glioblastoma. In fact, some studies have suggested that the combination of anti-PD-1 and anti-VEGF therapies may actually reduce the efficacy of each drug individually. Therefore, based on the current understanding of their mechanisms of action and available data, the drug combination is predicted to be non-synergistic in the treatment of glioblastoma.

- Drug combo: Nivolumab, Ipilimumab
- Targets: PD-1, CTLA-4
- Disease: advanced, pretreated, immunotherapy-naive squamous NSCLC
- Inferred synergy: Yes
- (False positive)

Explanation of prediction

- Nivolumab and Ipilimumab are monoclonal antibodies that function as checkpoint inhibitors by blocking the PD-1 and CTLA-4 pathways, respectively.
- Blocking PD-1 allows T cells to remain active and proliferate, increasing their ability to recognize and attack cancer cells.
- Blocking CTLA-4 allows T cells to become more effective in recognizing and attacking cancer cells by reducing the suppression of T-cell responses by antigen-presenting cells.

- The combination of PD-1 and CTLA-4 blockade can lead to a more robust and durable immune response compared to monotherapy with either drug alone.
- By targeting both PD-1 and CTLA-4, the drugs can enhance the immune system's response against cancer cells.

- Drug combo: Cediranib, Olaparib
- Targets: VEGFR, PARP
- Disease: high-grade serous and endometrioid ovarian cancer
- Inferred synergy: Yes
- (True positive)

Explanation of prediction

- The predicted synergy between Cediranib and olaparib in treating high-grade serous and endometrioid ovarian cancer is based on preclinical studies suggesting synergy between VEGFR and PARP inhibitors in other cancer types.
- Cediranib inhibits the activity of VEGFR, which is involved in angiogenesis and tumor growth. Olaparib, on the other hand, inhibits the PARP enzyme, which plays a role in DNA repair. By inhibiting both angiogenesis and DNA repair, the combination of Cediranib and olaparib may be more effective in killing cancer cells than either drug alone.
- Olaparib can sensitize cancer cells to the cytotoxic effects of Cediranib by inhibiting DNA repair mechanisms, making the cells more susceptible to the apoptotic effects of Cediranib.
- High-grade serous and endometrioid ovarian cancer often have mutations in genes such as BRCA1 and BRCA2, which are key targets for PARP inhibitors. These mutations result in a deficiency in DNA repair mechanisms, making cancer cells more susceptible to the cytotoxic effects of PARP inhibitors. Cediranib, by inhibiting angiogenesis, can help reduce the tumor size and improve the delivery of olaparib to the cancer cells, enhancing its efficacy.

- Drug combo: pembrolizumab, Docetaxel
- Targets: PD-1, tubulin
- Disease: Metastatic Castration-Resistant Prostate Cancer (mCRPC)
- Inferred synergy: No
- (True negative)

Explanation of prediction

- Pembrolizumab is an immune checkpoint inhibitor that blocks the interaction between PD-1 and its ligands, thereby releasing the brakes on the immune system and allowing T cells to attack cancer cells.
- Docetaxel, on the other hand, is a taxane chemotherapeutic agent that interferes with the stability of microtubules, leading to cell cycle arrest and apoptosis. While both drugs

have been used in the treatment of various cancers, including mCRPC, their mechanisms of action do not directly contribute to a synergistic effect.

- In fact, the use of immunotherapies like pembrolizumab and chemotherapies like docetaxel can sometimes have opposing effects, with chemotherapy potentially reducing the effectiveness of immunotherapy by damaging healthy cells and suppressing the immune response

- Drug combo: favezelimab, Pembrolizumab
- Targets: LAG3, PD-1
- Disease: Colorectal (PD-L1 positive MSS)
- Inferred synergy: Yes
- (False positive)

Explanation of prediction

- The synergistic effect of combining favezelimab (targeting LAG3) and pembrolizumab (targeting PD-1) in treating colorectal (PD-L1 positive MSS) cancer is predicted based on preclinical studies suggesting enhanced immune response when blocking both LAG3 and PD-1 checkpoints.
- LAG3 and PD-1 are co-inhibitory proteins that play important roles in regulating T-cell function and tumor immune evasion.
- By targeting both LAG3 and PD-1, the combination therapy is expected to result in more effective T-cell activation and proliferation, leading to a stronger antitumor response.
- This synergistic effect is thought to occur through the simultaneous release of inhibitory signals and the enhancement of T-cell activation.
